# SAR11 Cells Rely on Enzyme Multifunctionality to Transport and Metabolize a Range of Polyamine Compounds

**DOI:** 10.1101/2021.05.13.444117

**Authors:** Stephen E. Noell, Gregory E. Barrell, Christopher Suffridge, Jeff Morré, Kevin P. Gable, Jason R. Graff, Brian J. VerWey, Ferdi L. Hellweger, Stephen J. Giovannoni

**Author notes:** Address correspondence to Stephen J. Giovannoni.

## Abstract

In the ocean surface layer and cell culture, the polyamine transport protein PotD of SAR11 bacteria is often one of the most abundant proteins detected. Polyamines are organic cations produced by all living organisms and are thought to be an important component of dissolved organic matter (DOM) produced in planktonic ecosystems. We hypothesized that SAR11 cells transport and metabolize multiple polyamines and use them as sources of carbon and nitrogen. Metabolic footprinting and fingerprinting were used to measure the uptake of five polyamine compounds (putrescine, cadaverine, agmatine, norspermidine, and spermidine) in two SAR11 strains that represent the majority of SAR11 cells in the surface ocean environment, *Ca.* Pelagibacter st. HTCC7211 and *C.* P. ubique st. HTCC1062. Both strains transported all five polyamines and concentrated them to micromolar or millimolar intracellular concentrations. Both strains could use most of the polyamines to meet their nitrogen requirements, but we did not find evidence of use as carbon sources. We propose potABCD transports cadaverine, agmatine, and norspermidine, in addition to its usual substrates of spermidine and putrescine, and that spermidine synthase, speE, is reversible, catalyzing the breakdown of spermidine and norspermidine, in addition to its usual biosynthetic role. These findings provide support for the hypothesis that enzyme multifunctionality enables streamlined cells in planktonic ecosystems to increase the range of DOM compounds they oxidize.

**Importance:** Genome streamlining in SAR11 bacterioplankton has resulted in a small repertoire of genes, yet paradoxically they consume a substantial fraction of primary production in the oceans. Enzyme multifunctionality is hypothesized to be an adaptation that increases the range of organic compounds oxidized by cells in environments where selection favors genome minimization. We provide experimental support for this hypothesis by demonstrating that SAR11 cells use multiple polyamine compounds and propose that a small set of multifunctional genes catalyze this metabolism. We also report polyamine uptake rates can exceed metabolism, resulting in high intracellular concentrations of these nitrogen-rich compounds and an increase in cell size. Increases in cytoplasmic solute concentrations during transient episodes of high nutrient exposure has previously been observed in SAR11 cells and may be a feature of their strategy for maximizing the share of labile DOM acquired when in competition with other cell types.

## Introduction

Polyamines are low molecular weight organic polycations that are ubiquitous in living organisms. They play a role in stabilizing DNA, RNA, and proteins, are required for cell growth, and have been implicated in biofilm formation (1–3). Polyamine compounds and concentrations vary between cell types and can depend on nutrient status, temperature, and salinity (4). Polyamines are found at low nanomolar concentrations in the coastal and open ocean, reaching maximal concentrations of 30 nM during algal blooms, but typically are around 1 nM (5–7). Polyamines from the environment are metabolized by bacteria as nitrogen and carbon sources at rates similar to those of dissolved free amino acids and supply up to 14% of bacterial nitrogen demand in coastal regions. (8–10).

Putrescine (PUT) and spermidine (SPD), the most abundant polyamines in the ocean, are typically 3–5 nM in the environment, but spermine, cadaverine (CAD), and norspermidine (NSD) have been detected at lower levels (6, 10, 11). Several other polyamines, such as 1,3-diaminopropane (DAP), agmatine (AGM), homospermidine (HSD), spermine, and larger, more complex polyamines are known to be produced and/or metabolized by cells from all domains of life (3, 4, 9, 12, 13). Metabolic pathways for common polyamines are shown in Figure 1.

**Figure 1.**
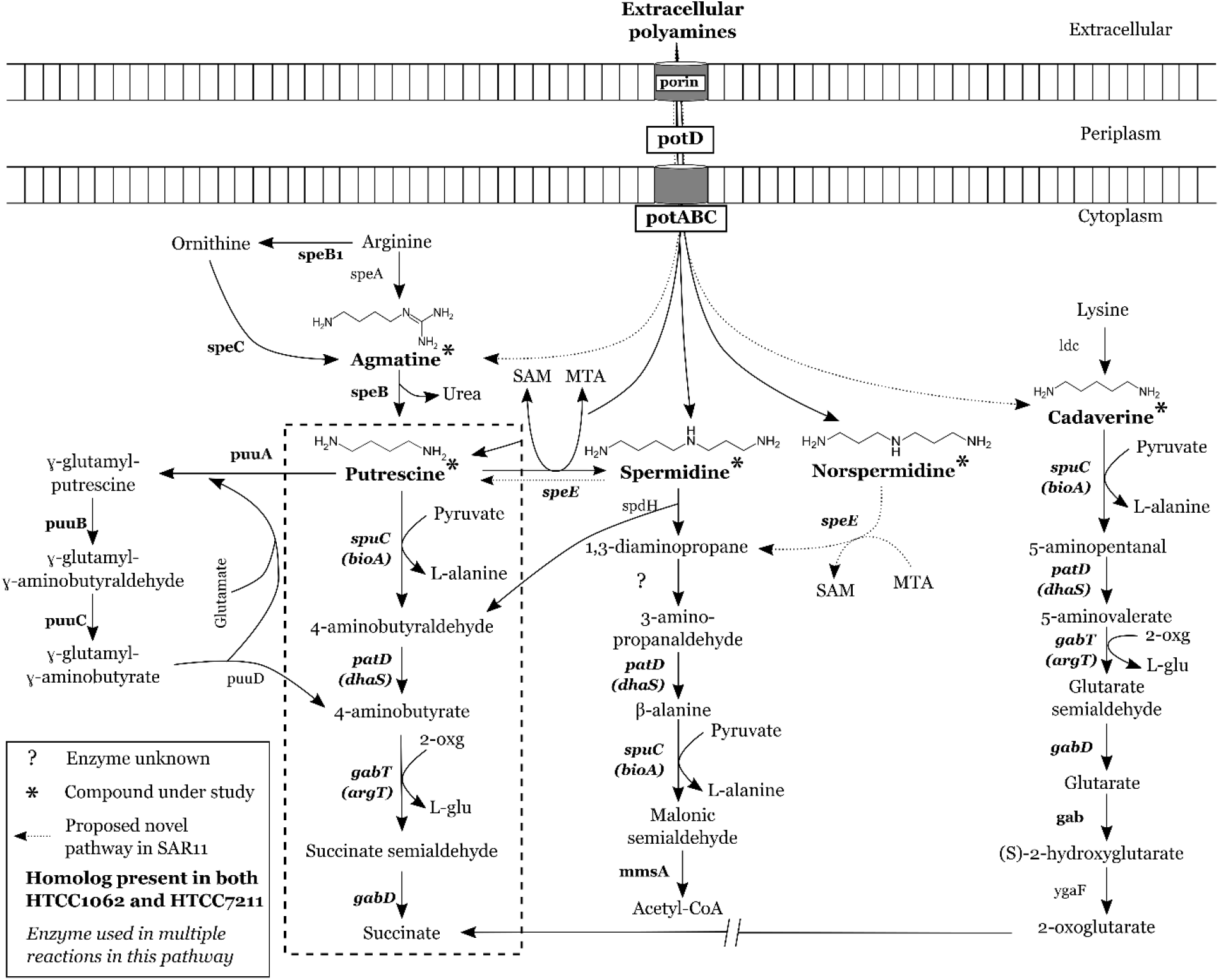
Polyamine compound metabolism in SAR11. Common pathways for polyamine metabolism in bacteria are shown. The compounds under study are marked with asterisks, with enzymes listed in bold if both strains of SAR11 in this study, HTCC1062 and HTCC7211, have homologs to the enzyme, or in plain text if in neither. A question mark indicates the enzyme is unknown. The enzyme name is in italics if it is used for multiple reactions in this metabolic system. The dashed box encompasses the pathway thought to be used by SAR11 cells for PUT metabolism based on previous studies. By-products of reactions where NH_3_ groups are transferred are included to show the flow of N. Gene names for SAR11 homologs, where different from canonical gene names are listed below the canonical name in parentheses. potC: spermidine/putrescine ABC transporter, permease; potB: permease; potD: SBP; potA: ATP-binding protein; speB1: arginase; speC: lysine/ornithine decarboxylase; speB: agmatinase; speE: spermidine/spermine synthase; puuA: Gamma-glutamylputrescine synthetase; puuB: Gamma-glutamylputrescine oxidoreductase; puuC: NADP/NAD-dependent aldehyde dehydrogenase; puuD: Gamma-glutamyl-gamma-aminobutyrate hydrolase; spuC: putrescine-pyruvate aminotransferase; kauB: 4-aminobutyraldehyde dehydrogenase; gabT: acetylornithine aminotransferase; gabD: succinate-semialdehyde dehydrogenase; spdH: spermidine dehydrogenase; malonate-semialdehyde dehydrogenase; gab: glutarate 2-hydroxylase; ygaF: L-2-hydroxyglutarate dehydrogenase. SAM: S-adenosylmethionine. MTA: 5’- methylthioadenosine. 2-oxg: 2-oxoglutarate. L-glu: L-glutamate

SAR11 alphaproteobacteria make up the majority of bacteria in the ocean (14). SAR11 cells primarily utilize labile, low molecular weight molecules (15). They pack their relatively large periplasmic space (16) with large numbers of ABC transporter substrate-binding proteins (SBPs) (17, 18), increasing the encounter rate and binding of substrate molecules with SBPs, resulting in high whole-cell uptake affinities (19, 20). Recent modeling work has extended this observation, suggesting this strategy may contribute to the slow growth rates of SAR11 cells (21). SAR11 bacteria evolved minimal genomes in response to streamlining selection, which favors efficient use of resources in nutrient-limited ecosystems (22). Enzyme multifunctionality has been hypothesized to reduce gene content in streamlined cells and has been confirmed for the SAR11 glycine betaine transporter (20).

SAR11 cells produce large numbers of PotD, the SBP involved in polyamine transport, both in cultures and the environment, making it the most highly expressed transporter for N-related compounds by SAR11 cells (17, 18, 23). N-limited cultures of SAR11 strain HTCC1062, a member of the cold, high-latitude Group Ia.1 ecotype, did not upregulate genes for polyamine transport or metabolism, except for an enzyme implicated in PUT and CAD metabolism (24), but genes involved in the metabolism and transport of other organic N sources were upregulated (24). Incubation experiments with natural seawater communities provided evidence that SAR11 cells may sometimes respond to additions of polyamines PUT and SPD - transcripts for SAR11 genes involved in polyamine metabolism increased in the first hour of incubation and accounted for over a quarter of all transcripts (25). In other experiments with PUT and SPD amendments to seawater, it was observed that SAR11 cell abundance did not change during a 48-hour period in response to PUT and SPD addition (26, 27); oligotrophs frequently decrease in relative abundance in incubation experiments due to their slow growth rates, while copiotrophs increase rapidly due to their fast growth rates under high nutrient conditions used in incubation experiments (15).

In this study we used targeted metabolic footprinting and fingerprinting to examine the types and amounts of polyamines taken up and metabolized by two SAR11 strains. Both strains of SAR11 used in this study come from the Ia subgroup: *Ca.* Pelagibacter ubique HTCC1062 belongs to the cold, high-latitude Group Ia.1 ecotype and *Ca.* P. st. HTCC7211 is from the equatorial, warm water Ia.3 ecotype (15). Targeted metabolic footprinting uses LC-MS/MS to measure changes in the concentrations of specific metabolites dissolved in spent culture media (28), while fingerprinting quantifies the concentrations of targeted metabolites within cells (29). We hypothesized that SAR11 cells would use polyamines as N sources and that polyamines might supply these cells with metabolic carbon via a branch of metabolism that proceeds through pyruvate (25, 30).

## Results

### Footprinting and Fingerprinting Experiments

We focused on five polyamine compounds (Table 1): putrescine (PUT), cadaverine (CAD), agmatine (AGM), norspermidine (NSD), and spermidine (SPD). These compounds were picked either for their prevalence in the environment and in bacteria cells or for their role as precursors to other polyamine compounds (4, 6, 31, 32). These compounds also showed the best recovery in solid-phase extraction and were amenable to simultaneous quantification by LC-MS/MS. We used polyamine concentrations 10-100X ambient environmental concentrations, similar to what would be found in nutrient patches (33), as has been done previously (20, 25). In preliminary experiments, we found that high polyamine concentrations inhibited growth; HTCC1062 growth was inhibited when all polyamines were added together at individual concentrations above 500 nM, while HTCC7211 growth was inhibited at concentrations above 250 nM. We chose 500 nM for HTCC1062 and 250 nM for HTCC7211, the highest concentrations that did not significantly inhibit growth, for further experiments (Figure S1(A-B)).

**Table 1.**
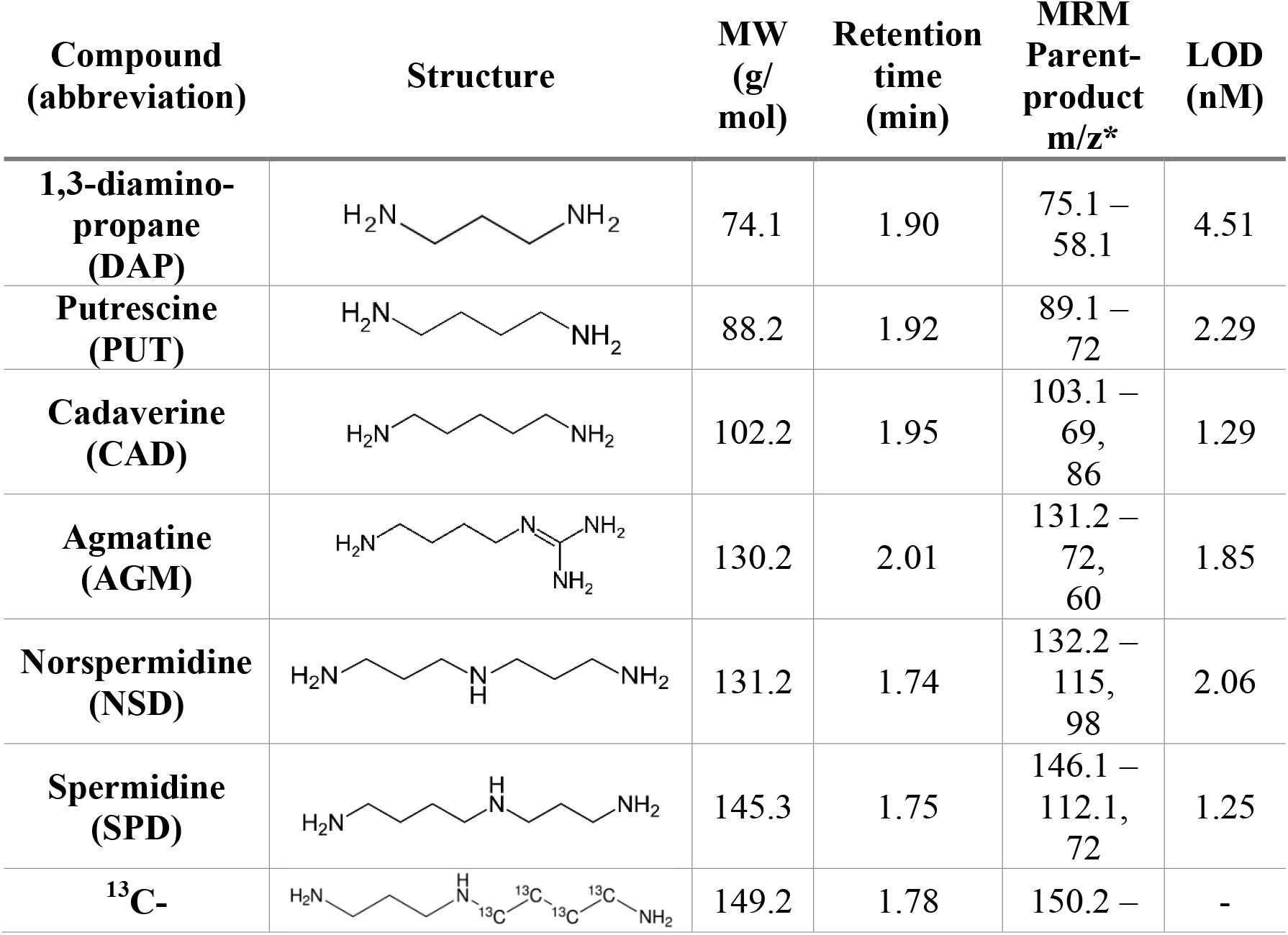
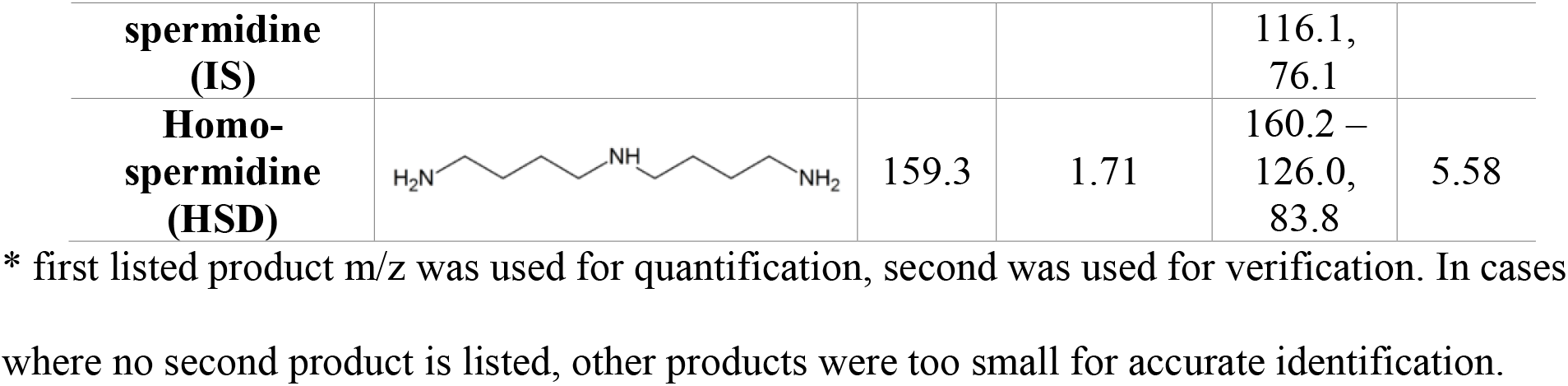
List of compounds used in project as well as LC-MS/MS parameters. Instrumental limit of detection (LOD) for the LC-MS/MS was calculated as three times the standard deviation of the lowest detectable concentration of standards used (5 nM). The actual LOD of samples varied based on how much the samples were concentrated, but was generally 2 – 1000X lower. A collision energy of 20 eV was used for all analytes except 1,3-diaminopropane, which had a CE of 10 eV, and homospermidine, which had a CE of 30 eV.

| Compound<br>(abbreviation) | Structure | MW<br>(g/<br>mol) | Retention<br>time<br>(min) | MRM<br>Parent-<br>product<br>m/z* | LOD<br>(nM) |
| --- | --- | --- | --- | --- | --- |
| **1,3-diamino-**<br>**propane**<br>**(DAP)** |  | 74.1 | 1.90 | 75.1 –<br>58.1 | 4.51 |
| **Putrescine**<br>**(PUT)** |  | 88.2 | 1.92 | 89.1 –<br>72 | 2.29 |
| **Cadaverine**<br>**(CAD)** |  | 102.2 | 1.95 | 103.1 –<br>69,<br>86 | 1.29 |
| **Agmatine**<br>**(AGM)** |  | 130.2 | 2.01 | 131.2 –<br>72,<br>60 | 1.85 |
| **Norspermidine**<br>**(NSD)** |  | 131.2 | 1.74 | 132.2 –<br>115,<br>98 | 2.06 |
| **Spermidine**<br>**(SPD)** |  | 145.3 | 1.75 | 146.1 –<br>112.1,<br>72 | 1.25 |
| **<sup>13</sup>C-** |  | 149.2 | 1.78 | 150.2 – | - |
| **spermidine (IS)** |  |  |  | 116.1, 76.1 |  |
| **Homo-spermidine (HSD)** |  | 159.3 | 1.71 | 160.2 – 126.0, 83.8 | 5.58 |

\* first listed product m/z was used for quantification, second was used for verification. In cases where no second product is listed, other products were too small for accurate identification.

The five polyamines were added to SAR11 cultures under nutrient replete conditions to measure uptake and oxidation of these compounds. Cultures were grown to late exponential phase before harvesting; growth rates were slightly lower with polyamines added: 0.44 compared to 0.46 d^−1^ for HTCC1062 and 0.50 compared to 0.60 d^−1^ for HTCC7211 with and without polyamines (Figure S2(C-D)). For both strains, average intracellular levels of all five polyamine compounds were significantly greater in the experimental treatment (polyamines added) compared to the negative control (no polyamines added), except for SPD in HTCC1062, which had non-significant higher levels in experimental cultures (Figure 2A, C; p-values in Table 2). When the intracellular levels are converted to intracellular concentrations using a cell volume of 0.03 μm^3^ for HTCC1062 (16) and 0.04 μm^3^ for HTCC7211 (34), it is apparent that the cells are concentrating all compounds into intracellular concentrations greater than their environment (Table 2). The intracellular concentrations in the experimental treatment for HTCC7211 were much higher than in HTCC1062, especially SPD, which was 40X higher.

**Table 2.** Intracellular concentrations of polyamine compounds in SAR11 fingerprinting experiments (see Figure 2), calculated using cell volumes of 0.03 and 0.04 μm^3^ for HTCC1062 and HTCC7211, respectively. Listed p-values are for a one-sided t-test comparing the + Polyamines treatment to the negative control treatment.

| Compound | HTCC1062 intracellular concentration (mM) |  |  | HTCC7211 intracellular concentration (mM) |  |  |
| --- | --- | --- | --- | --- | --- | --- |
|  | - Control | + Polyamines | p-value | - Control | + Polyamines | p-value |
| **Putrescine** | 0.016 ± 0.0067 | 0.093 ± 0.023 | 0.003 | 0.006 ± 0.0003 | 4.03 ± 1.86 | 0.03 |
| **Cadaverine** | 0 | 0.0015 ± 0.001 | 0.03 | 0 | 0.153 ± 0.031 | 0.0002 |
| **Agmatine** | 0 | 0.003 ± 0.0021 | 0.03 | 0 | 1.46 ± 0.94 | 0.02 |
| **Nor-spermidine** | 0 | 0.0038 ± 0.0016 | 0.01 | 0 | 1.02 ± 0.49 | 0.003 |
| **Spermidine** | 0.328 ± 0.127 | 0.48 ± 0.13 | 0.19 | 0.012 ± 0.0027 | 19.1 ± 9.63 | 0.01 |

**Figure 2.**
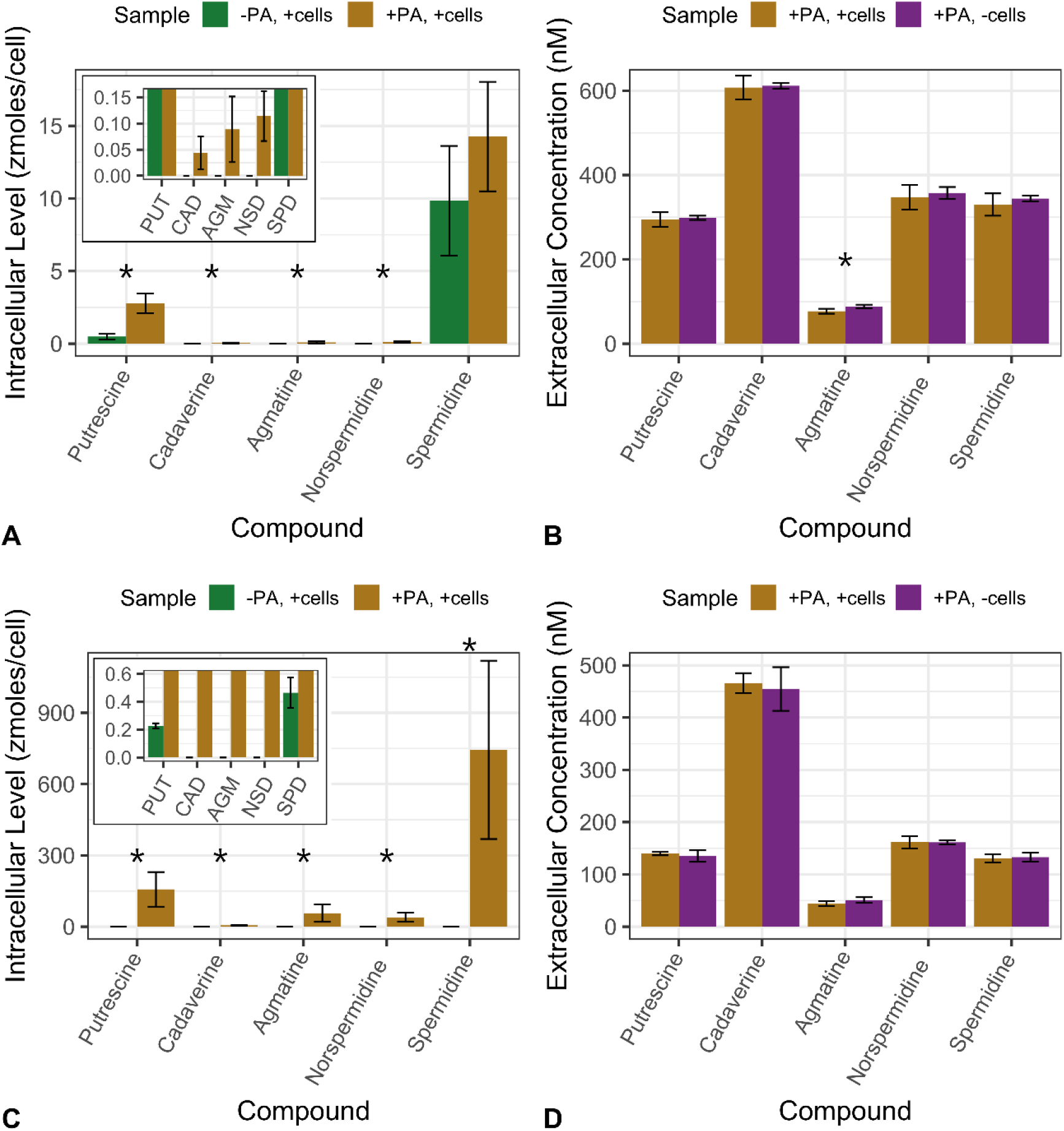
Results of footprinting/fingerprinting experiments with (A, B) Ca. P. ubique HTCC1062 and (C, D) Ca. P. st. HTCC7211 with five polyamine compounds. (A, C): intracellular concentrations of polyamine compounds in cultures with no polyamines added or with 500 nM or 250 nM final concentration of each polyamine compound added for HTCC1062 or HTCC7211, respectively. (B, D): extracellular concentrations of polyamine compounds in cultures with 500 nM or 250 nM final concentration of each polyamine compound added for HTCC1062 or HTCC7211, respectively. Insets in (A) and (C) show compounds that had very low levels. Error bars are standard deviation of quadruplicate cultures. * indicates significant difference (p < 0.05, one-tailed t-test) between the two treatments; for (A) and (C), indicates significantly higher level of that compound in experimental treatment compared to the negative control; for (B) and (D), indicates significantly lower concentration of that compound in experimental treatment compared to the no cells control; for p-values, see Table 2.

In the extracellular fractions of both strains, there were no significant differences between the experimental treatment and the no cell control (polyamines added, no cells), except for AGM in HTCC1062 (p-value of 0.04, one-sided t-test) (Figure 2B, D). In HTCC1062, all five compounds were lower in concentration in the experimental treatment compared to the no cell control (Figure 2B). For HTCC7211, all five compounds were at similar concentrations between the two treatments, except for AGM, which was lower in the experimental treatment (p-value of 0.06, one-sided t-test).

### Flow Cytometry Experiment

Based on the very high intracellular polyamine concentrations measured in HTCC7211 cells in experimental treatments, we postulated that cells would change in size to accommodate the influx of polyamines. To test this, we used flow cytometry to monitor the forward scatter (FSC), a proxy for cell size, of nutrient-replete HTCC7211 cultures exposed to either no polyamines (control) or 250 nM of each polyamine added at the beginning of growth (early addition) or after 4 days of growth (late addition). On average, both experimental treatments had higher FSC than the control (Figure 3B). Mean FSC of the early addition cultures was consistently higher than the control across all measured time points (Figure 3B). Mean FSC of late addition cultures was similar to early addition cultures 4 hours after addition of polyamines to late addition cultures; it then decreased to control levels at day 7, finally increasing above the control after day 11 (Figure 3A).

**Figure 3.**
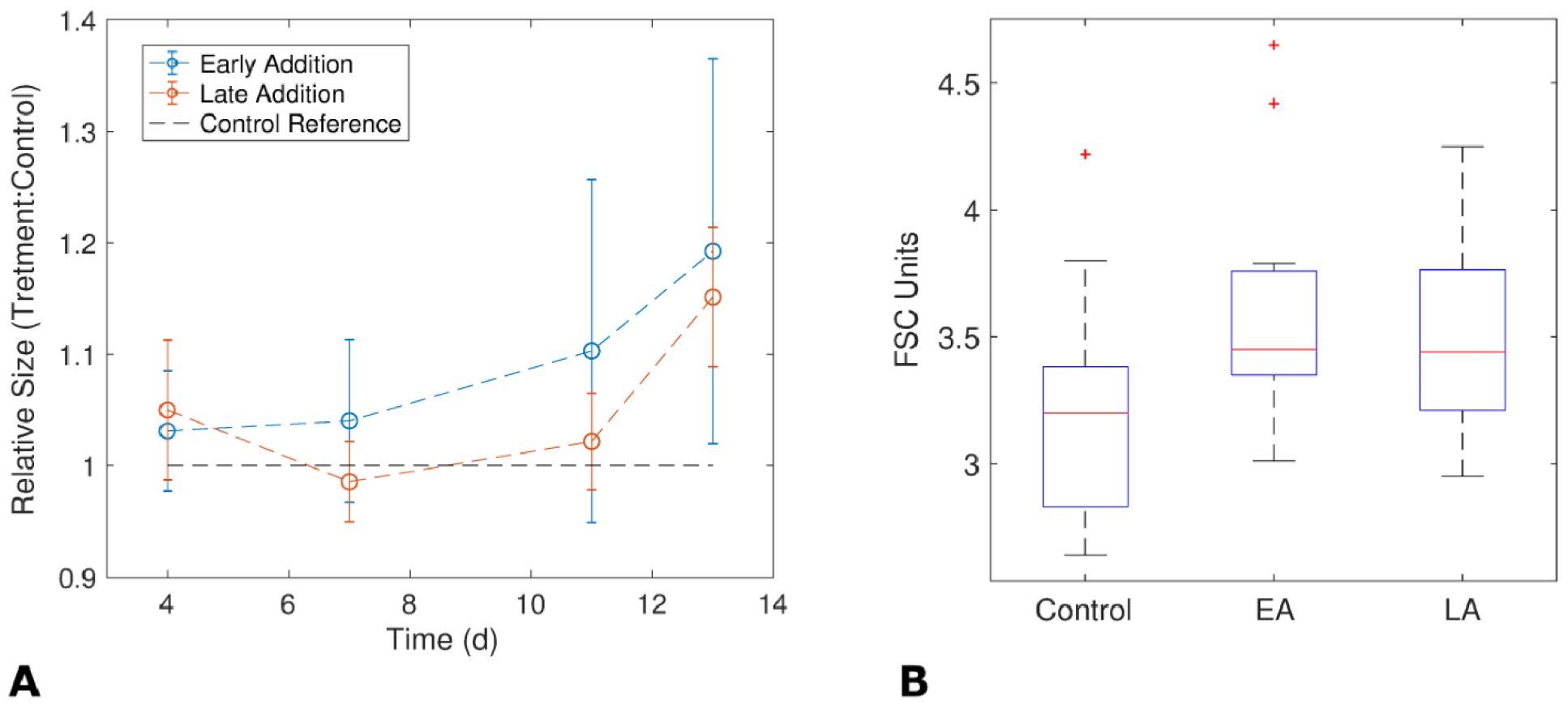
Changes in cell size (FSC) were monitored using flow cytometry in response to addition of 250 nM polyamines either at the beginning of the experiment (early addition) or after 4 days of growth (late addition). (A) Mean FSC from experimental treatments after normalization to the mean of the control treatment cultures (no polyamines added). (B) Mean FSC (proxy for cell size) for the three treatments over the time course. Error bars are the standard deviation of quadruplicate replicates.

### Carbon Substitution Experiments

Growth experiments were used to examine whether the five polyamine compounds could substitute for two unusual growth requirements of SAR11 cells: pyruvate, or related compounds which lead to a branch of SAR11 metabolism that includes the biosynthesis of alanine, and glycine or related compounds, required for another branch of SAR11 metabolism that includes glycine synthesis. The five polyamine compounds were added together at final concentrations of 250 nM each as a replacement for either pyruvate or glycine, and the growth of the cultures compared to negative control treatments with either no pyruvate or no glycine added. With both strains, experimental treatments with added polyamines achieved higher maximum cell densities and more rapid growth rates than negative controls, but the differences were not significant, indicating that these compounds did not substitute for glycine or pyruvate (Table 3).

**Table 3.** Growth rates and maximum cell densities for growth experiments with SAR11 cultures to determine if polyamines could substitute for pyruvate or glycine in their growth. All five polyamine compounds were added in at 250 nM final concentration each in place of either pyruvate or glycine and compared to negative controls with either no pyruvate or glycine added. Pyruvate was added at 100 μM and glycine at 50 μM; 10 μM methionine and vitamins were added to all cultures in artificial seawater medium (see Methods).

| **SAR11 Strain** | **Treatment** | **Growth rate**<br>**(day<sup>-1</sup>)**<br>**± stdev** | **Maximum cell**<br>**density**<br>**(cells/mL)**<br>**± stdev** |
| --- | --- | --- | --- |
| **HTCC1062** | + Control (+ glycine,<br>+ pyruvate) | 0.33 ± 0.02 | 3.27E7 ±<br>5.72E6 |
|  | - Control (- glycine) | 0.21 ± 0.003 | 4.19E6 ±<br>7.63E5 |
|  | Experimental (- glycine,<br>+ polyamines) | 0.23 ± 0.02 | 5.30E6 ±<br>1.66E6 |
|  | - Control (- pyruvate) | 0.14 ± 0.03 | 1.25E6 ±<br>4.65E5 |
|  | Experimental (- pyruvate,<br>+ polyamines) | 0.15 ± 0.01 | 1.36E6 ±<br>2.22E5 |
| **HTCC7211** | + Control (+ glycine,<br>+ pyruvate) | 0.55 ± 0.00 | 1.48E8 ±<br>1.98E7 |
|  | - Control (- glycine) | -0.11 ± 0.01 | 6.64E4 ±<br>1.17E3 |
|  | Experimental (- glycine,<br>+ polyamines) | -0.12 ± 0.1 | 7.57E4 ±<br>2.37E4 |
|  | - Control (- pyruvate) | 0.19 ± 0.02 | 1.04E6 ±<br>2.55E5 |
|  | Experimental (- pyruvate,<br>+ polyamines) | 0.22 ± 0.03 | 1.76E6 ±<br>3.41E5 |

### Nitrogen Substitution Growth Experiments

Additional growth experiments were used to determine if polyamine compounds could serve as nitrogen (N) sources. To eliminate other organic sources of N, oxaloacetate and dimethylsulfoniopropionate (DMSP) were substituted for glycine and methionine in a modified artificial seawater medium (24). These substitutions resulted in ~100X lower maximum cell densities due to oxaloacetate being a weak glycine substitute. The five polyamine compounds were added to SAR11 cultures grown in the modified ASW media either all together at a final concentration of 150 nM for each compound, or individually at concentrations that provided equivalent amounts of N (245 nM N). Maximum cell densities were compared to a negative control with no added N (−C), a positive control with excess N in the form of ammonium sulfate (+C, excess N), and a positive control with equimolar N in the form of ammonium sulfate (+C, equimolar N) (Figure 4).

**Figure 4.**
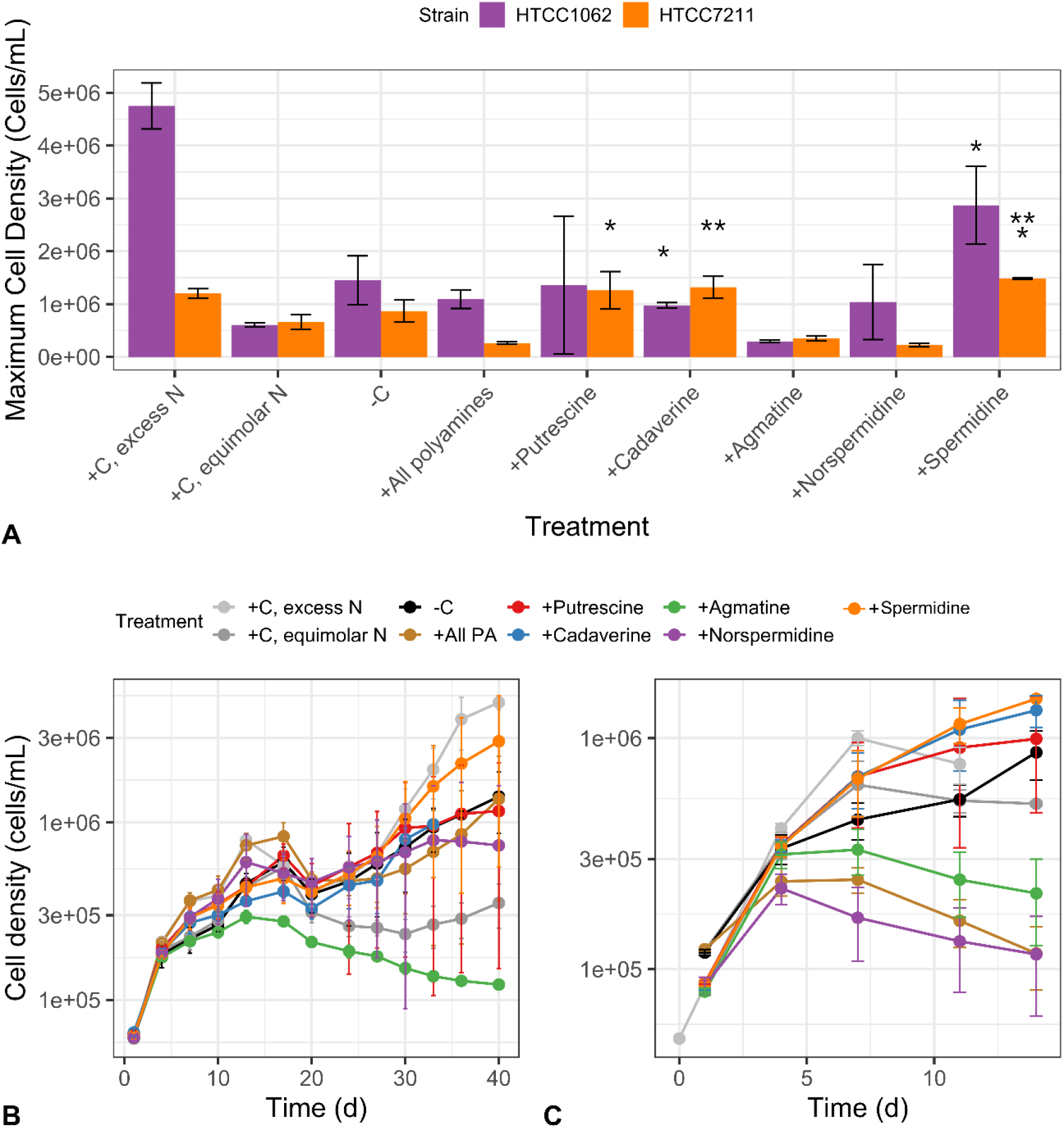
Several polyamine compounds are used by SAR11 cells as a nitrogen (N) source, as indicated by higher maximum cell densities during growth experiments compared to control cultures. SAR11 cells (either HTCC1062 or HTCC7211) were grown on a modified ASM recipe (see Methods) without any N source (−C). Positive control cultures were grown with N in the form of (NH_4_)_2_SO4, with either 400 μM (+C, excess N) or with equimolar N equal to the amount of N (245 nM N) added in the polyamine treatments (+C, equimolar N). Experimental treatments had equal amounts of N (245 nM N) added in the form of either all five polyamine compounds together, final concentration of 150 nM for each compound (+All polyamines), or with individual polyamine compounds. Cultures were started from cultures that had been grown to late exponential phase on the same media without any N, to eliminate any carryover from previous growth. Error bars represent the standard deviation of triplicate samples. *: significantly higher maximum cell density compared to the +C, equimolar N treatment (one-sided t-test, p < 0.05); **: significantly higher maximum cell density compared to both +C, equimolar N and -C treatments (one-sided t-test, p < 0.05); ***: significantly higher maximum cell density compared to all three control treatments (one-sided t-test, p < 0.05). For HTCC7211, results from two separate experiments are shown together.

For HTCC1062, addition of two compounds, CAD and SPD, resulted in significantly higher maximum cell densities than the equimolar positive control (maximum cell densities and p-values in Table S2) (Figure 4). Addition of NSD, PUT, and all polyamines combined resulted in higher, albeit non-significant, maximum cell densities than the equimolar positive control. Only addition of SPD resulted in a higher cell density than the negative control, but the difference was not significant. For HTCC7211, addition of SPD resulted in a significantly higher maximum cell density than all three controls. Cultures to which CAD was added had significantly higher maximum cell densities than the equimolar control and the negative control. Cultures with PUT added had a significantly higher maximum cell density than only the equimolar control. In both strains, the addition of AGM resulted in lower maximum cell densities than any control treatments; in HTCC7211, the treatments with NSD and with all polyamines combined also had lower maximum cell densities than any controls. Interestingly, diauxic growth was observed in HTCC1062, with an early peak around 14 d and a larger peak later around 40 d (Figure 4), which was not observed in HTCC7211.

### Metabolic Pathways

Figure 1 shows genes for polyamine metabolism for the two SAR11 strains used in this study, overlayed on common pathways for polyamine metabolism (8, 12, 30, 35–37). In both strains, AGM is postulated to be converted by agmatinase (*speB*) to PUT, which is catabolized by the transamination pathway, since neither strain encodes the final enzyme in the γ-glutamylation pathway, and the transamination pathway was upregulated in SAR11 cells in response to PUT addition (25). CAD is likely metabolized to succinate via the lysine degradation pathway. Many genes involved in polyamine metabolism are known to be multifunctional in other cell types, as we also predict in SAR11 (Figure 1, Figure S2).

In metabolic reconstruction from genome sequences (38), we found that neither SAR11 strain encoded a clear pathway of SPD or NSD metabolism, although both compounds were taken up from the medium and metabolized (Figure 2, Figure 4). Microbial enzymes responsible for NSD metabolism have not been extensively characterized. Several *Vibrio* strains produce and metabolize NSD via carboxynorspermidine (39), but the necessary enzymes are lacking in SAR11. Neither SAR11 strain has homologs for the canonical genes responsible for SPD metabolism: SPD dehydrogenase (s*pdH*), which cleaves SPD to produce 1,3-diaminopropane (DAP) and 4-aminobutanal, and SPD acetyltransferase, which converts SPD to acetylspermidine, a less toxic version of SPD. We speculate that either the SPD synthase enzyme, SpeE, is bi-directional, producing PUT from SPD, or there is another, unknown enzyme capable of cleaving SPD. SpeE is not known to be bi-directional in other bacteria (40). A possible candidate enzyme for SPD metabolism was discovered during metabolic reconstruction: *dys2*, a putative deoxyhypusine synthase (*dhs*) gene (Figure S). *Dys2* is highly conserved in these and other SAR11 genomes between two genes involved in PUT formation, *speC* and *speB*. In prokaryotes, *dhs* usually acts as a homospermidine (HSD) synthase, a promiscuous enzyme capable of acting on multiple polyamines in addition to its native function of producing HSD from PUT, instead of the function it performs in eukaryotes and archaea of post-translationally modifying elongation factor 5 (EF5) while cleaving SPD (41, 42).

To differentiate between these two alternative routes of SPD metabolism and identify pathways of NSD metabolism in SAR11, we used fingerprinting to search for possible by-products of SPD and NSD metabolism, including DAP and HSD. If SAR11 cells use the reverse SPD synthase reaction to metabolize SPD, the products would be PUT and S-adenosylmethionine, while the by-products of the Dys2 enzyme, if it is a HSD synthase, would primarily be HSD and DAP. We compared polyamine levels in SAR11 cells grown either without any polyamines or with either 500 nM SPD or NSD (Figure 5). As expected, in both strains, the treatments with SPD or NSD added had higher levels of that compound in the respective treatment (one-sided t-test, p-values of 0.02 and 0.003 for +SPD and +NSD vs. the negative control, respectively; p-values were 0.06 and 0.2 for HTCC1062), indicating an uptake of those two compounds by both strains.

**Figure 5.**
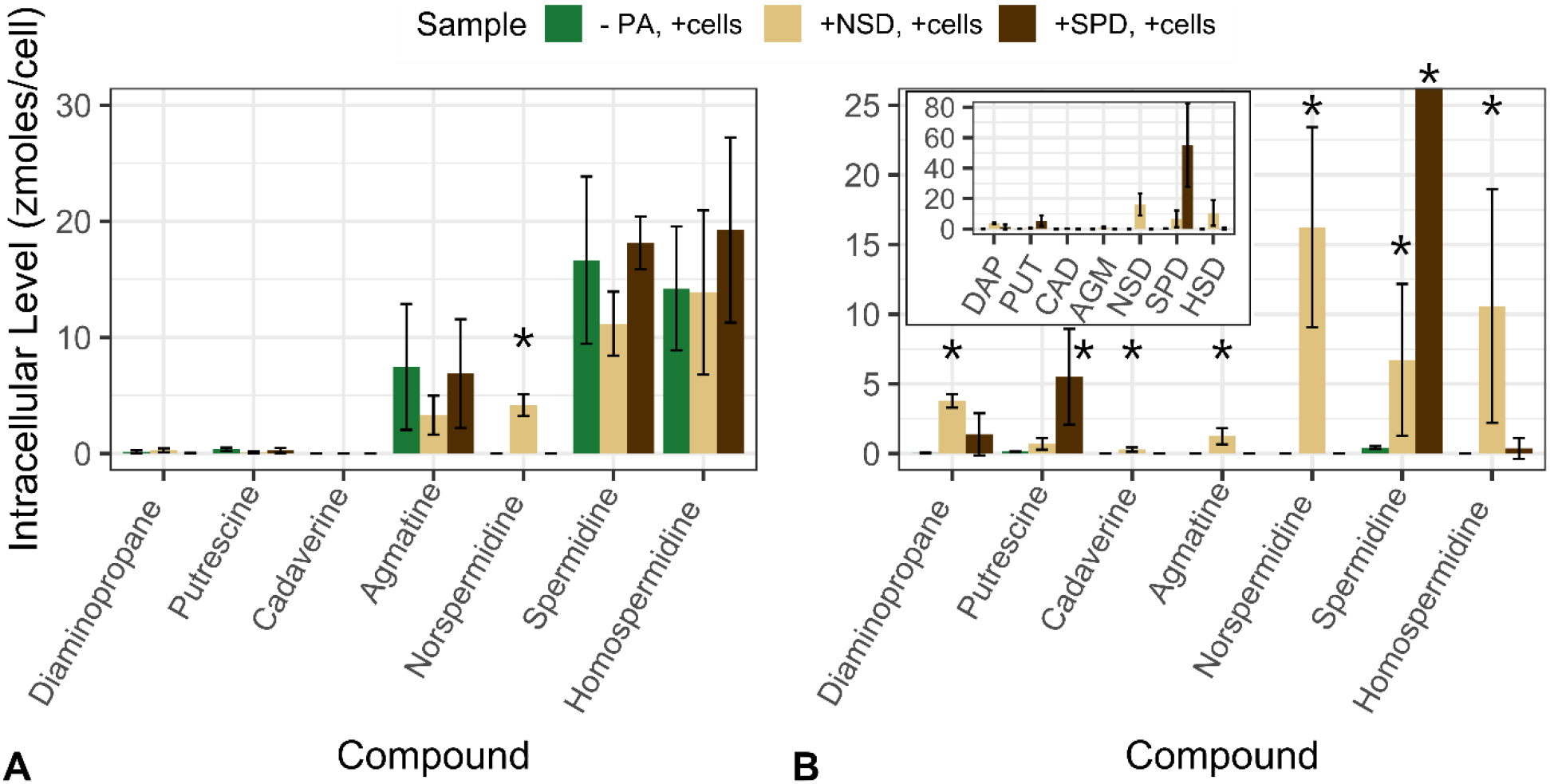
Intracellular polyamine levels indicating pathways for norspermidine (NSD) and spermidine (SPD) metabolism in (A) HTCC1062 and (B) HTCC7211. Cultures were grown in nutrient replete conditions either without any added polyamines, with 500 nM NSD, or with 500 nM SPD added. Cells were harvested at late exponential phase and intracellular polyamines extracted as described in the Methods section. * indicates a significantly higher level of that compound compared to the control treatment with no polyamines added (one-sided t-test, p < 0.05). Error bars are the standard deviation of quadruplicate cultures.

In HTCC7211, the +SPD cultures also had significantly higher levels of PUT than the negative control (one-sided t-test, p-value of 0.05) but only a slight, non-significant increase in DAP (Figure 5B). In the +NSD cultures, there were significantly higher levels of DAP than the negative control (one-sided t-test, p-value of 0.001). There was also a significant increase in SPD and HSD (one-sided t-test, p-values of 0.05 and 0.04) in +NSD cultures compared to the negative control. In HTCC1062, there were no differences in the levels of any other compounds in the +NSD or +SPD treatments compared to the negative control, aside from SPD and NSD themselves and a non-significant increase in DAP in the +NSD treatment (Figure 5A).

## Discussion

SAR11 cells devote much of their cellular structure and energy to transport functions to compete for resources in the world’s nutrient-limited oceans (17, 18, 20). One of the primary transporters expressed by SAR11 cells, PotABCD, transports polyamines, yet little is known about the types and amounts of polyamines used by SAR11, the role of polyamines in SAR11 growth, and the pathways used by SAR11 for polyamine metabolism. Here, we show that SAR11 cells transported and metabolized all five polyamine compounds tested and concentrated polyamines to μM and even mM intracellular concentrations. Cells increased in size while concentrating polyamines, probably as a consequence of osmosis and increasing pressure inside the cells. We also show that SAR11 cells primarily use polyamines as a N source and propose metabolic pathways for SPD and NSD, two compounds for which catabolic pathways in marine bacteria are uncertain.

The growth of SAR11 cultures was found to be inhibited by high polyamine concentrations (Figures S1, 3). Polyamines are known to be toxic to bacteria when added to media at high concentrations (generally mM range), but the mechanism is not known (37, 43, 44). SAR11 cells often lack transcriptional regulation for carbon oxidation functions (15) and previous work indicated they do not upregulate metabolic enzymes for polyamines when N limited (24). The growth inhibition observed at high polyamine concentrations might be due to adverse effects of the buildup of polyamine compounds inside the cells. Similar results previously have been observed in cells experiencing metabolic pathway saturation (45). Intracellular concentrations of polyamines were higher in HTCC7211 than in HTCC1062 and HTCC7211 was also more susceptible to growth inhibition by polyamines, indicating these might be linked (Figure S1(A-B), Table 2).

### Uptake of Polyamine Compounds by SAR11 Cells

Organisms produce intracellular polyamines for a variety of cellular processes including stabilization of DNA, RNA, and proteins (3). The native polyamines produced by these strains (SPD and HSD in HTCC1062 and SPD alone in HTCC7211 (Figure 2, Figure 5)) are consistent with past reports that alphaproteobacteria primarily make PUT, SPD, or HSD as polyamines (46). AGM and PUT, intermediates in the synthesis of SPD and HSD (Figure 1), were also detected at low levels. The concentrations of native polyamines measured in SAR11 cells are similar to those measured in various phytoplankton (4), but ~10X lower than found in other alphaproteobacteria (46).

Both strains transported all five polyamines in excess of metabolism, causing higher (10 – 1000X higher) intracellular concentrations in experimental treatments relative to negative controls (Figure 2A, C). Intracellular polyamines reached μM concentrations in HTCC1062 and mM concentrations in HTCC7211, 20 mM in the case of SPD in HTCC7211 (Table 2). In response to the large influx of polyamine compounds, HTCC7211 cells increased in size within hours, as indicated by the increase in FSC in the late addition treatment cultures at the first measured time point (Figure 3).

Substrate uptake in excess of metabolic rate has been observed previously in SAR11 cells with the osmolyte dimethylsulfoniopropionate (DMSP) (47). A parallel phenomenon, termed “luxury uptake,” has been described in phytoplankton that take up phosphorous and nitrogen in excess of their requirements and store them in organic forms for later use (48, 49). However, the excess substrate uptake we observed does not fit the canonical definition of luxury uptake, since the amount of N from the stored polyamines in SAR11 cells was far less than their requirement for a cell division: concentrated polyamines inside SAR11 cells in the experimental treatment only made up 0.005% for HTCC1062 and 0.27% for HTCC7211 of the cellular N quota, estimated at 0.11 fmol N/cell (34).

Theoretically, the excess uptake such as what we observed enhances the ability of cells to exploit nutrient patches, giving them a cache of nutrients to process subsequently after exiting a nutrient patch. Moreover, given that SAR11 cells are by far the most abundant cell type in the ocean, luxury uptake by SAR11 cells could be a population-level strategy that lowers ambient nutrient concentrations to levels where SAR11 cells are more competitive for transport, effectively taking nutrients off the table. Past theoretical work has shown that superior competitors in patchy environments lower average nutrient concentrations (50). Our findings may stimulate further research aimed at understanding whether these cellular behaviors apply to other substrate compounds used by SAR11 cells, whether similar behavior is exhibited by other oligotrophs, and whether the imbalance between transport and metabolism we observed occurs in natural populations and plays an adaptive role by allowing transient storage of exometabolites inside cells.

Although HTCC1062 and HTCC7211 transported all five compounds from the media, there were few observable depletions of extracellular concentrations for any polyamines with either strain because the intracellular polyamine pools were small relative to the surrounding volume (Figure 2C, D). The observed accumulation of intracellular polyamines was estimated to result in pmolar drawdowns of the dissolved polyamine pool, which in most cases was less than the precision of our measurements (Table S3). An exception was the accumulation of 239 pmoles of intracellular SPD in HTCC7211, which should have produced a measurable depletion of SPD in the medium, yet no significant reduction was observed (standard deviation of 83.3 pmoles). This observation suggests that HTCC7211 cells used other transported polyamines to synthesize SPD. To support this interpretation, HTCC7211 cells given only SPD had 10X lower SPD levels than when given all five compounds (compare Figure 2C and Figure 5B). It appears that transported polyamines are converted intracellularly to SPD, which then accumulates in HTCC7211.

Our analysis indicates the PotABCD transport system in SAR11 is responsible for transporting the five polyamines we tested, given the structural similarity between these compounds and the absence of other candidates for polyamine transport functions. Both strains of SAR11 lack homologs for CAD, NSD, or AGM transporters found in other bacteria. In *Vibrio cholerae*, NSD is transported by a *potABCD* homolog (51), but CAD and AGM have not previously been identified as substrates for *potABCD*. In *E. coli*, PotABCD is primarily a PUT/SPD/SPM transporter (52), which could help explain the higher accumulation of these compounds in SAR11 cells, even when accounting for native PUT and SPD production. This is the second multifunctional transporter identified experimentally in SAR11 cells, in addition to the ABC transporter for glycine betaine, which was shown to transport seven different substrates (20). It is likely that other ABC transporters in SAR11 cells are also multifunctional, given the use of a wide variety of amino acids and carboxylic acids by SAR11 cells (53–55).

Interestingly, there was a large difference between the two SAR11 strains in the amount of polyamines taken up. Polyamine concentrations in HTCC7211 were 40 – 500X higher than HTCC1062, despite HTCC7211 being exposed to 2X lower concentration of polyamines. This difference cannot be explained entirely by cell size; HTCC7211 cells are only ~1.3X larger, as measured by C content, than HTCC1062 (34). The differential could be because the HTCC7211 transport system has a higher V_max_ for polyamine transport than HTCC1062, due either to the properties of the proteins themselves (one of the two permease proteins involved in polyamine transport, PotB, is only 82% identical), differing abundance of transport proteins, or the cytoarchitecture of the cells. There were no major differences between the two strains in presence/absence of polyamine metabolic genes, nor in the location of those genes (Figure S2). One possible ecological explanation for the difference between these two strains, if they are typical of the ecotypes they represent, is that HTCC1062, a member of the primarily coastal subclade of SAR11, may have been influenced by selection that limits toxic buildups of polyamines at the higher polyamine concentrations found in coastal regions. HTCC7211, a member of a primarily open ocean subclade of SAR11, would rarely experience the high polyamine concentrations found in coastal regions and so might not experience selection to limit intracellular buildups.

### Use of Polyamines by SAR11 Cells

SAR11 cells have unique growth requirements, needing a reduced sulfur source (e.g. methionine or methane thiol), a glycine source, specific vitamins, and a carbon source that can serve as precursor to alanine (usually pyruvate) (54). Most of the tested polyamine compounds are predicted to be metabolized to succinate, a TCA cycle intermediate (Figure 1). In previous work, succinate was shown to not substitute for pyruvate in HTCC1062, in accord with our experimental findings (56). This does not rule out the use of polyamines as an energy source, however. Other small compounds have been found previously to be used by SAR11 cells only as an energy source and not as a pyruvate substitute (20, 55). Polyamines are also required for a variety of other cellular processes, and it is likely that SAR11 cells used the supplied polyamines in those processes in addition to metabolizing them.

In both strains of SAR11 tested, several polyamine compounds (SPD, CAD, and PUT) were able to be used as a N source, with SPD supporting the highest maximum cell density of any of the polyamines (Figure 4A). NSD does not appear to be a N substitute for either strain of SAR11 at the NSD concentration tested, but it is transported (Figure 5). The use of multiple polyamines as a N source is consistent with previous reports that SAR11 cells use a variety of organic N-containing compounds as N sources (24).

Interestingly, several compounds (AGM and NSD) were inhibitory to SAR11 growth under N limiting conditions (Figure 4). One potential cause for the AGM inhibition is the by-product of AGM degradation by the agmatinase enzyme, urea. Neither SAR11 strain encodes a urease (57). We speculate that the influx of AGM causes a build-up of inhibitory urea in cells, a process that is known to occur in oligotrophs due to metabolic pathway saturation (15, 45). This does not, however, preclude AGM from also being used as a N source by the cells at environmental AGM concentrations.

Surprisingly, all HTCC1062 cultures, including controls, exhibited diauxic growth during N-substitution experiments, while HTCC7211 cultures, under similar conditions, did not. Diauxic growth is generally observed when cells switch from using one source of nutrients to another. However, the cultures used to start these experiments were acclimated to the same medium (without N) prior to the experiment starting. Previously, diauxic growth was observed in HTCC7211 grown on alternate P sources, which was attributed to the switch from using inorganic P to organic P sources (58). Another unexpected observation, found in both strains and across several repetitions, was that the equimolar positive control always had a lower maximum cell density than the negative control with no N added.

### Metabolic pathways for Spermidine and Norspermidine

SPD metabolism has been observed in marine bacteria without a spermidine dehydrogenase gene (*spdH*) (9, 11, 25). It has been speculated that the enzyme that synthesizes SPD from PUT, SpeE, is bi-directional, although this activity was not confirmed experimentally (9). In HTCC7211, it appears that SPD is primarily metabolized via the reverse SPD synthase reaction, not via the Dys2 enzyme, as no significant increase in HSD was detected, while an increase in PUT was observed (Figure 5B). The SpeE enzyme in SAR11 previously has been found to be the result of a gene fusion event and is highly multifunctional, displaying high biosynthetic activity with multiple polyamines substrates, in addition to this enzyme’s commonly predicted substrate, PUT (59). Our data suggest that this enzyme is not only multifunctional in substrate range, but also in its ability to carry out catalytic reactions in reverse of its usual biosynthetic activities. The results from HTCC1062 on SPD metabolism are not as clear, as no other differences between the +SPD treatment and the negative control were observed aside from an increase in SPD (Figure 5A).

We propose that NSD is being metabolized in HTCC7211 by the enzyme SpeE, similar to SPD, since we observed increased levels of DAP in the NSD treatment and the SpeE enzyme in SAR11 is known to have a wide substrate rage (Figure 5B). With these data, we cannot rule out the Dys2 enzyme metabolizing NSD. In HTCC7211, there was also an increase in SPD and HSD in the +NSD treatment (Figure 5B). The Dys2 enzyme in SAR11 cells may be responsible for producing SPD and HSD from the excess NSD and DAP, since *dhs* homologs in bacteria are known to produce SPD from PUT and DAP, in addition to producing HSD from PUT (42, 60, 61). It appears that the *dys2* gene in both SAR11 strains is not primarily acting as a HSD synthase, since there was relatively low production of HSD under any condition, in contrast to other prokaryotes with a HSD synthase gene where HSD is the sole polyamine present (60, 61).

We evaluated the hypothesis that SPD synthase might be catalyzing reactions that are the reverse of its ordinary action of synthesizing polyamines. We explored thermodynamic models that predicted the energies of the compounds in the primary reaction catalyzed by SPD synthase (forming SPD from PUT; Figure S3), without considering entropy terms (Text S1, Table S4). The estimated ∆E value for the total reaction was positive (14.32 kcal/mol) when water was used as the proton acceptor (it is expected that ΔH° values will be quite similar to ΔE values) (Table S5). More favorable acceptors (e.g., imidazole) easily yield negative ∆E values of −23.49 kcal/mol (Table S5). Our findings suggest that the direction of this reaction is easily tunable by including proton carriers of varying strengths. For this calculation, we used standard conditions and did not consider the very high intracellular concentrations of polyamines we observed experimentally, which might further drive this reaction to reverse its normal biosynthetic function.

## Conclusion

Some properties of plankton cells that are important to understanding and modeling their behavior in natural ecosystems can only be measured by experimentation. Recently, we demonstrated very low whole-cell affinities and multifunctionality in the osmolyte transport system of cultured SAR11 cells. We attributed these competitively advantageous cell properties to synergism between kinetic features of the glycine betaine transporter ProXYZ and unusual aspects of SAR11 cell architecture, notably their small size and large periplasm packed with abundant substrate binding proteins (20). Here, we investigate SAR11 metabolism of polyamines, which are transported into cells by the highly abundant SAR11 transporter system PotABCD. We find this system is also multifunctional, and that the two SAR11 strains metabolized a variety of polyamines, which served the cells as N sources. We cannot rule out polyamines, which we predict to be metabolized to succinate, being used by SAR11 as carbon sources, but our findings do not show that polyamines are a major source of carbon for these cells. In previous work, we have shown that several C1 compounds are used by SAR11 as energy sources via tetrahydrofolate-mediated oxidation but not as a source of carbon for biomass production (55), which may be the case with polyamines. Our data strongly support the hypothesis that SAR11 use many polyamines via a simplified system of few enzymes and a single transporter. They mainly use these compounds as an N source and perhaps to supplement their intracellular polyamine pool, potentially important adaptations in N-limited marine systems.

Polyamine transport rates exceeded metabolic rates, leading to mM intracellular polyamine accumulations and an increase in cell size over a period of hours. We propose that SAR11 cells use the multifunctional enzyme spermidine synthase, SpeE, in reversible reactions that can both produce SPD and catabolize SPD and NSD. The findings we report indicate enzyme multifunctionality expands the range of DOM compounds these cells harvest, which may partially explain how these cells attain high success in competition for DOM resources. Our findings also support previous observations which indicated SAR11 cells concentrate some metabolites during pulses of availability, metabolizing them subsequently (47). In principle this cell behavior could increase the success of SAR11 cells in competition for nutrient patches, but further experimental work and modeling are needed to evaluate this hypothesis. In any case, the properties of cells that we uncovered here are neither typical nor trivial - they change our understanding of how competition for DOM resources has led to the emergence of specialized cell types and will likely inform future experimental research and modeling aimed at understanding cell evolution and the ocean carbon cycle.

## Methods

### SAR11 Growth and Washing

The protocol for SAR11 growth conditions, cell counts, and washing has been previously reported (20). Briefly, *Candidatus* Pelagibacter sp. HTCC7211 and *Ca.* P. ubique HTCC1062 were grown on artificial seawater (ASW) amended with 100 μM pyruvate, 50 μM glycine, 10 μM methionine, and SAR11-specific vitamins at either 25 or 16 °C (HTCC7211 and HTCC1062, respectively) in 12 h light/dark (54, 56). For testing polyamines as a nitrogen (N) source, a modified ASW was used without inorganic N (ammonium sulfate). This modified ASW was amended with 100 μM pyruvate, 50 μM oxaloacetate (instead of glycine), 1 μM dimethylsulfoniopropionate (DMSP; instead of methionine), and vitamins, since both glycine and methionine provide potential organic N sources (24).

In the growth experiments testing polyamines as a N source, there were two positive controls: a N-replete positive control, with 400 μM ammonium sulfate added, and an equimolar N positive control, containing the same stoichiometric ammonium sulfate concentration as the experimental cultures with organic N sources (polyamines). For growth experiments, cell counts were taken at least twice a week and growth rates and maximum cell densities calculated as described previously (54). For growth experiments, a starting cell density of 1E5 cells/mL was used; a density of 5E4 cells/mL was used in experiments using the modified ASW, as cells grew to a lower maximum cell density in this medium. For the experiments testing polyamines as a N source, cultures were started from cultures grown without N to late exponential phase to exhaust any carryover N.

For footprinting/fingerprinting experiments, cells were harvested using centrifugation (Beckman-Coulter J2-21) at 10°C at 30,000 g for 90 min. Cell pellets were re-suspended in TE buffer and pelleted again (Beckman-Coulter Ultracentrifuge) at 12°C at 48,000 g for 60 minutes (20).

### Footprinting and Fingerprinting Experiments

To measure the identity and quantity of polyamine compounds used by SAR11 cells, a panel of five common polyamine compounds was added to SAR11 cultures at the beginning of growth under normal growth conditions. The polyamine compounds used were putrescine (PUT), cadaverine (CAD), agmatine (AGM), norspermidine (NSD), and spermidine (SPD). Each polyamine compound was added at a final concentration of 500 nM for HTCC1062 or 250 nM for HTCC7211. 250 nM was used for HTCC7211 as they were found to be inhibited in growth at 500 nM (Figure S1 (A-B)). Concentrations used were about 100X ambient polyamine concentrations for several reasons: to allow for accurate quantification of polyamines; to match the high densities of cells used, which were also about 100X average cell densities in the ocean; and to match previous amendment studies with polyamines and other, similar compounds (25, 27, 62). A negative control treatment with SAR11 cells but no polyamines added was used to measure native polyamine production and background polyamine concentrations in the ASW. A no-cell control with polyamines and no SAR11 cells was also included to account for degradation of polyamines, extraction efficiency during SPE, and carryover polyamines in the intracellular measurements. All treatments were done in quadruplicate. Samples were taken for cell counts twice a week; samples for extracellular (footprint) and intracellular (fingerprint) polyamine measurements were taken in late exponential phase.

The metabolic pathways of two of the polyamine compounds, SPD and NSD, are not clear in SAR11 cells. Thus, further fingerprinting experiments were carried out with SPD and NSD added separately to cultures at 500 nM. Several possible metabolic intermediates and byproducts were measured in these cells and compared to a negative control with no polyamines added.

### Intracellular Polyamine Extraction

Intracellular polyamines were extracted from SAR11 cells using a method adapted from targeted marine metabolomics studies (63). Cells were pelleted and washed in TE Buffer as described above. The supernatant was then completely removed and the cell pellet was re-suspended in 10 μL of fresh TE buffer. Volume was determined via an analytical balance. 1 μL of re-suspended cells was removed and diluted 1:200 in TE buffer to determine cellular abundance for per-cell normalization. Cells were lysed by adding 100 μL of cold methanol (MeOH; LC-MS grade) to the cell suspension, followed by an addition of 300 μL of cold 1 M acetic acid (63). Cell lysis and polyamine extraction was completed by shaking samples on high for 5 min followed by rest in ice for 1 min to avoid overheating, repeated three times. A liquid phase extraction was used to remove hydrophobic components of the cellular matrix: 400 μL of chloroform (LC-MS grade) was added and the samples shaken for 1 min, followed by centrifugation for 5 min at 5,000 RPM to achieve phase separation. The aqueous layer containing the polyamines was transferred into a new tube and the organic layer discarded. The resulting sample was concentrated via drying under a nitrogen stream at 30°C. The dried samples were re-suspended in 30 μL of 50:50 1 M acetic acid:acetonitrile (64), weighed, and analyzed as described below. When 125 nM standards of each compound were extracted using this method, minimal degradation was observed, with over 60% recovery (Table S1). Because no intracellular polyamine standard reference exists, reported intracellular measurements in this paper were not corrected for recovery efficiency.

### Extracellular SPE Extraction

Polyamines dissolved in the culture media were extracted using a solid phase extraction (SPE) as described previously (64). 10 mL of the supernatant from the initial centrifugation of cultures (described above) was used for each sample. Polyamines were extracted via gravity alone (nominal flow rate of 0.07 mL/min) onto a 1000 mg Bond Elut-C_18_ SPE column (3 mL, Agilent) pre-conditioned with methanol and bicarbonate buffer at pH 12. Salts were removed from the column by washing three times with 1 mL of 0.1 M borate buffer, pH 12. Polyamines were eluted into cryovials with three washes of equal volumes of 1 M acetic acid and acetonitrile, final volume 5 mL. Greater than 85% recovery for CAD, PUT, NSD, and SPD was observed, while AGM had 48% recovery (Table S1), perhaps because of the high pKa of AGM (~12) compared to the other four compounds (~10). Artificial seawater (blanks) extracted using this method only had minimal concentrations (less than 4 nM) of NSD and SPD and none of the other three compounds (data not shown). Measured concentrations were not corrected for extraction recovery and so values reported in this paper are conservative.

### LC-MS/MS Analysis

Quantification of polyamines was carried out using an Applied Biosystems 4000 Q-Trap triple quadrupole mass spectrometer with an ESI interface, coupled to a Shimadzu LC-20AD liquid chromatograph (LC-MS/MS). Applied Biosystems Analyst and ABSciex Multiquant software packages were used for instrument operation and quantification respectively. A Phenyl-3, 150 × 4.6 mm 5 μm HPLC column (GL Sciences) was used for chromatographic separations, using a 2.0 μm pre-filter as a guard column (Optimize Technologies). The sample rack was cooled to 10°C to prevent degradation of polyamines. The column temperature was maintained at 40°C. HPLC mobile phases were MS grade water (Fisher) with 0.1% formic acid and MS grade acetonitrile (Fisher) with 0.1% formic acid. A 10-minute binary gradient with a flow rate of 0.8 mL/min was used. The initial concentration of 3% acetonitrile ramped to 30% acetonitrile in 5 minutes. The column then re-equilibrated at 3% acetonitrile for 5 minutes. The ESI Source used a spray voltage of 5200 V and source temperature of 600 C. Sheath gas pressure was 50 PSI and auxiliary gas pressure was 40 PSI. The mass spec was run in positive ion mode. Compound-specific MRM parameters, column retention times, and limits of detection are presented in Table 1. The instrumental limits of detection (LOD; Table 1) were calculated as three times the standard deviation of six runs of the lowest detectable standard (5 nM). Sample injection volume was 10 μL and samples were analyzed in triplicate. Samples and standards were all analyzed in a 50:50 acetonitrile:acetic acid mix. Samples were randomized prior to analysis. ^13^C-spermidine (Spermidine-(butyl-^13^C4) trihydrochloride, Sigma Aldrich) was added as an internal standard (IS) for quantification and to compensate for matrix effects. Compound concentrations that are listed as zero in figures and tables in this paper were below the LOD. LCMS analysis was conducted at the Oregon State University Mass Spectrometry Core Facility. Data analysis was conducted in the R software environment (R Core Team, 2015), and all figures were created using the Ggplot2 software package for R (R Core Team; Wickham, 2016).

### Flow Cytometry Analysis

Flow cytometry was used to monitor changes in cell size and morphology in HTCC7211 cultures in response to exposure to polyamines. Three treatments were used, all with cells grown under nutrient replete conditions, with quadruplicate cultures for each treatment; cultures were started at a cell density of 5E4 cells/mL. The negative control had no polyamines added; in one experimental treatment (early addition), cultures were started with 250 nM of each polyamine added to the media; in the other experimental treatment (late addition),250 nM of each polyamine was added after 4 days of growth, when they had passed 5E5 cells/mL density. Samples were taken twice a week until they reached early stationary phase. Cells were stained with SYBR green I prior to analysis on a Becton Dickinson Influx Cell Sorter (BD ICS). The BD ICS was equipped with a 488 nm laser and detectors for forward scatter (FSC), side scatter (SSC) and fluorescence at 530 nm, the emission wavelength of SYBR Green I, and 692 nm. Prior to sample analysis, the instrument was aligned according to manufacture specification. See (65, 66) for additional information on instrument specifications, alignment, and calibration procedures. Data were recorded for >50,000 cells per replicate per treatment. Flow cytometry files were analyzed using FlowJo (v.10.7.1) and cells were identified by their fluorescence in the 530 nm channel. The mean for each replicate for each detector channel was recorded and used in calculating the treatment means and standard deviations. Unfortunately, errors with the flow cytometer prevented observations of cells prior to day 4 of the experiment and data is restricted to day 4, 7, 11, and 13. Cell concentrations determined on the flow cytometer are presented in Figure S1(G).

### Computational Modeling

The reaction catalyzed by spermidine synthase (Figure S3) was computationally modeled to determine reaction energetics. Water and histidine were used separately as bases (B:). Amine nitrogens in putrescine and spermidine were fully protonated, but the nitrogens in the adenosyl fragment were left unprotonated (neutral). First, conformational spaces for compounds I and III were explored using Spartan’14 (67), with the MMFF force field (Figure S4). DFT studies were performed using Gaussian16 (68). The B3LYP functional (69–72) was employed using the cc-pVDZ basis set (73). An SCRF solvation model using water (74) was applied. All structures were optimized and showed only real vibrational frequencies. SCF energies with solvation correction were used as the primary measure of molecular energy. Reaction energies were estimated using the minimum energy conformer for each compound.

## Acknowledgments

This work was funded by the National Science Foundation grants OCE-1436865 and IOS-1838445. This work was supported, in part, by the Oregon State University Research Office. The content is solely the responsibility of the authors and does not necessarily represent the official views of the OSU Mass Spectrometry Center. The authors acknowledge the OSU Mass Spectrometry Center at Oregon State University and the specific institutional instrument grant for Applied Biosystems 4000 Qtrap: NIH # 1S10RR022589-01.

## Legends for Supplementary Material

Table S1 Recoveries for each of the polyamine compounds using the two extraction methods used. Extracellular recovery is the recovery percentage of standards extracted from artificial seawater (ASW) media using solid phase extraction (SPE). 500 nM standards of each compound were added to 10 mL ASW and extracted as described in the Methods. Reported values are the average and standard deviation of triplicate samples. For the intracellular recovery, 125 nM standards of each compound in 10 μL volume were extracted using the intracellular extraction method described in the Methods. Reported values are the average and standard deviation of duplicate samples.

Table S2 Maximum cell densities for SAR11 cultures grown with polyamine compounds as the sole nitrogen source with corresponding p-values. All p-values are from a one-sided t-test to assess whether the treatment had a significantly higher maximum cell density than the corresponding control. When an experimental treatment’s maximum cell density was lower than or equal to the corresponding control, a t-test was not conducted.

Table S3 Intracellular polyamine measurements from fingerprinting experiments (see Figure 2) fall within the error range of corresponding extracellular footprint measurements from the spent culture media. The intracellular polyamine levels in cells grown on polyamines were back calculated to mole values based on the number of cells in the culture when harvested. The standard deviation (stdev) for the extracellular measurements of polyamine compounds from the corresponding spent media were also back calculated to mole values for comparison. The only compound that clearly falls outside the error measurements is spermidine in HTCC7211.

Table S4 Computed energies for the compounds of interest in the spermidine synthase reaction (Figure S3). I: S-adenosyl-3-(methylsulfanyl)-propylamine; III: S-methyl-5’-thioadenosine.

Table S5 Estimated energies of reaction for the canonical spermidine synthase reaction (Figure S3) using either water or imidazole as the proton acceptor. It is expected that ΔH° values will be quite similar to ΔE values.

Figure S1 Growth experiments were carried out to determine the maximum concentration of polyamine compounds that can be added without negatively affecting growth of either (A) *Ca.* P. ubique HTCC1062 or (B) *Ca.* P. st. HTCC7211 under nutrient replete conditions (see main text, Methods for media used). Compounds were added at final concentrations of either 0 (control), 100, 250, 500, or 1000 nM for each compound. Error bars are standard deviation of triplicate samples. (C, D) Cell count data from the cultures used for footprinting experiments for either (C) HTCC1062 or (D) HTCC7211. The treatments with polyamines (PA) added had either 500 nM (HTCC1062) or 250 nM (HTCC7211) final concentration for each of the five polyamines added. Error bars are standard deviation of quadruplicate samples. (E, F) Cell count data from the cultures used in experiments to measure for the presence of 1,3-diaminopropane and homospermidine in either (E) HTCC1062 or (F) HTCC7211. The treatments with norspermidine (NSD) and spermidine (SPD) added had 500 nM final concentration of each added. Error bars are standard deviation of quadruplicate samples. (G) Cell count data from an experiment to monitor changes in cell size and morphology in response to polyamine addition in HTCC7211 cells using flow cytometry. Control: no polyamines added. Early addition: 250 nM of each polyamine compound was added at the beginning of growth. Late addition: 250 nM of each polyamine compound was added after 4 days of growth. Error bars are standard deviation of quadruplicate cultures.

Figure S2 Loci for genes involved in polyamine metabolism (marked with a *) in the two strains of SAR11 under study, HTCC1062 and HTCC7211. Gene abbreviations and corresponding enzyme names are listed here; gene abbreviations in parentheses represent the homologous gene. speB: agmatinase; ppiA: peptidylprolyl cis-trans isomerase precursor; fadB: 3-hydroxyacyl-CoA dehydrogenase; potC: spermidine/putrescine ABC transporter, permease; potB: permease; potD: SBP; potA: ATP-binding protein; prr: betaine-aldehyde dehydrogenase; gab: glutarate 2-hydroxylase; oocM: octopine transporter, permease; occQ: octopine transporter, permease; occt: octopine transporter, SBP; speE: spermidine/spermine synthase; pfs (mtnN): 5’-methylthioadenosine/S-adenosylhomocysteine nucleosidase; apt: adenine phosphoribosyltransferase; mtnA: methylthioribose-1-phosphate isomerase; mtnP: methylthioadenosine phosphorylase; speC: lysine/ornithine decarboxylase; dys2: deoxyhypusine synthase-like protein; speB1: arginase; yhhQ: uncharacterized ACR; rpmB: 50S ribosomal protein L28; pyrD: dihydroorotate dehydrogenase; lysS: lysine-tRNA ligase; aldH (kauB): 4-aminobutyraldehyde dehydrogenase; bioA (spuC): putrescine-pyruvate aminotransferase; ald: alanine dehydrogenase; argD (gabT): acetylornithine aminotransferase; argF: ornithine carbamoyltransferase; cycM: cytochrome C; kdsB: 3-deoxy-manno-octulosonate cytidylyltransferase; gabD: succinate-semialdehyde dehydrogenase.

Figure S3 Canonical reaction catalyzed by the spermidine synthase enzyme. B: is the base used in the thermodynamic calculations as the proton acceptor in the reaction.

Figure S4 Different conformations of the two more complex compounds in the spermidine synthase reaction. (A) “Compact” conformer of S-adenosyl-3-(methylsulfanyl)-propylamine; (B) “Extended” conformer of S-adenosyl-3-(methylsulfanyl)-propylamine. (C) “Compact” conformer of S-methyl-5’-thioadenosine. (D) “Extended” conformer of S-methyl-5’-thioadenosine.

